# Propagating patterns of activity across motor cortex facilitate movement initiation

**DOI:** 10.1101/549568

**Authors:** Karthikeyan Balasubramanian, Vasileios Papadourakis, Wei Liang, Kazutaka Takahashi, Matt Best, Aaron J. Suminski, Nicholas G. Hatsopoulos

## Abstract

Voluntary movement initiation involves the modulation of neurons in the primary motor cortex (M1) around movement onset. Yet, similar modulations of M1 activity occur during movement planning when no movement occurs. Here, we show that a sequential spatio-temporal pattern of excitability based on beta oscillation amplitude attenuation propagates across M1 prior to the initiation of reaching movements in one of two oppositely oriented directions along the rostro-caudal axis. Using spatiotemporal patterns of intracortical microstimulation, we find that reaction time increases significantly when stimulation is delivered against but not with the natural propagation orientation suggesting that movement initiation requires a precise recruitment pattern in M1. Functional connections among M1 units emerge at movement onset that are oriented along the same rostro-caudal axis but not during movement planning. Finally, we show that beta amplitude profiles can more accurately decode muscle activity when these patterns conform to the natural propagating patterns. These findings provide the first causal evidence that large-scale, spatially organized propagating patterns of cortical excitability and activity are behaviorally relevant and may be a necessary component of movement initiation.

## Introduction

The initiation of voluntary skeletal movements is often associated with large-scale modulations of neurons in the primary motor cortex (M1)^1,2^. It remains a mystery, however, why similar modulations in M1 neurons during movement preparation, visual observation of action, and motor imagery occur in the absence of movement ^3–7^. One possibility is that modulation is weaker during these conditions and fails to exceed non-linear, thresholds in downstream spinal circuitry, thus resulting in no movement^5,8,9^. Yet, preparatory activity as measured during an instructed-delay paradigm often equals or exceeds movement-related activity particularly in premotor cortex which is known to provide substantial projections to the spinal cord^10,11^. Another possibility is that there is an active inhibitory gating mechanism preventing movement initiation that is released when movement needs to begin^12^ as has been shown during REM sleep^13,14^. There is, however, no evidence that such a gate exists in the context of voluntary skeletomotor behaviors. A third possibility argues that the motor cortex resides in a very high dimensional state space that allows patterns of ensemble activity related to preparation to evolve in dimensions that do not generate movement (i.e. the “output null space”) in contrast to movement-relevant activity patterns that reside in an “output potent space”^15^. Aligned with this idea, we argue that a specific spatio-temporal pattern of M1 excitability and unit modulation is required for voluntary movement initiation.

Oscillations in the beta frequency range (i.e. 15-35 Hz) as measured by the local field potential (LFP) are prevalent in various motor structures including the cerebellum, basal ganglia, and cortical areas such as M1^16–18^. Immediately prior to movement onset, the amplitude of these beta oscillations attenuates (i.e. desynchronize) in M1 when single units begin modulating^19,20^ (Figure 1a). The degree of beta oscillation attenuation is correlated with the magnitude of motor-evoked potentials as measured with transcranial magnetic stimulation^21^. Furthermore, the timing of beta attenuation with respect to the go signal varies with reaction time but is fixed with respect to movement onset indicating that beta attenuation is a neural correlate of movement initiation (Figure 1b; Supplementary Figure 1; see ^22,23^).

**Figure 1.**
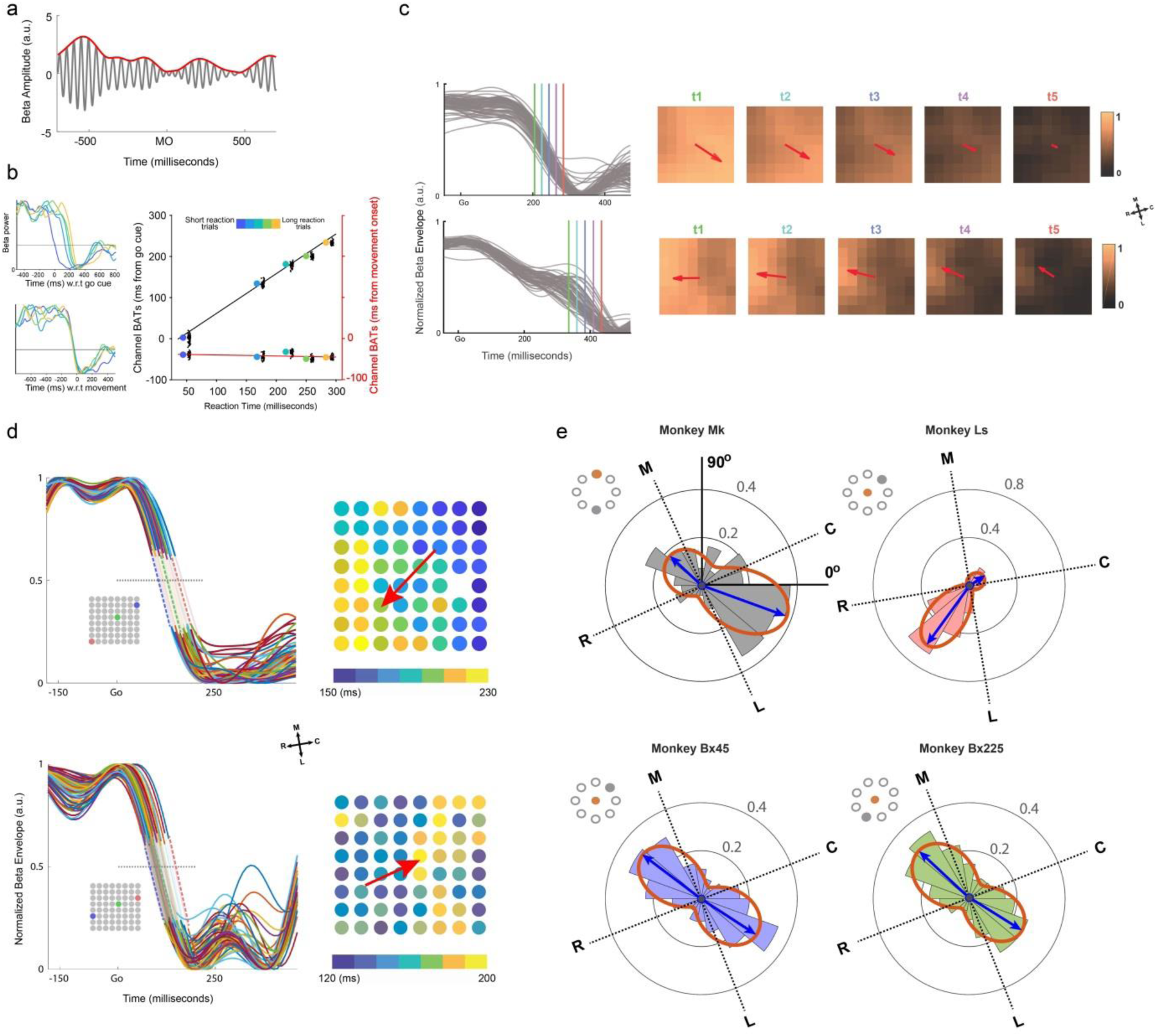
Single trial beta attenuation and their orientations. **a.** Beta frequency oscillations (gray) and amplitude envelope (red) on one trial from one electrode. MO-movement onset. **b.** Normalized beta envelopes averaged over different trial groups (trials sorted by reaction time, 50 trials per group) aligned on the GO cue (top left) or on movement onset (bottom left) from one session in Ls. (Right) Channel-averaged beta attenuation time regressed on mean reaction times (big colored dots) in each trial group with respect to the GO cue (black line, r^2^= 0.996, p=0.00018) or movement onset (red line, r^2^= 0.126, p=0.558). Black dots represent single channel values (offset by 10ms and jittered for clarity). **c.** Beta envelopes across channels from Monkey Ls for two trials (left) along with their corresponding spatial patterning (right) at different times (corresponding to vertical colored lines on left panel). Orientations of regression fits to the spatial patterning are denoted with arrows. **d.** Beta envelopes of two trials (left; inset shows spatial positions of three electrodes with early (blue), intermediate (green), and late (red) attenuation times) and their corresponding spatial maps of beta attenuation times (right) are shown for monkey Bx. Regression fits to the pattern show the beta attenuation orientation (BAO) directed rostrally or caudally (red arrows). **e.** Polar histograms show the distributions of BAOs across multiple single trials and recording sessions for three monkeys. A mixture of two von Mises functions was fit to the BAO distributions (red curves) to estimate the two direction modes of the BAO (blue arrows). Insets represent the initial hold (filled red) and target positions (filled gray) of the hand.

We have previously documented that trial-averaged, beta attenuation timing (BAT) does not occur synchronously across the upper limb area of M1 but rather forms a propagating sequence of cortical excitability^24^. Here, we extract single-trial beta amplitude attenuation profiles and find that attenuation propagates along one of two oppositely oriented directions along the rostro-caudal axis. By applying propagating patterns of subthreshold intracortical microstimulation (ICMS) prior to movement onset, we significantly delayed reaction time when ICMS patterns were incongruent with the natural propagating patterns. Using a network analysis on spiking activities of simultaneously recorded units, we observe that functional connections emerge immediately prior to movement onset that are oriented along the same rostro-caudal axis. Finally, by decoding muscle EMG activity from patterns of beta amplitude profiles and then applying spatial perturbations, we provide evidence that these natural propagating sequential patterns lead to normal recruitment of muscle activity.

## Results

### Single-trial propagating patterns of cortical excitability

Three rhesus macaque monkeys (Mk, Ls, and Bx) were trained on a simple reaction time task to perform planar reaching movements by moving a cursor to visual targets using a two-link robotic exoskeleton (BKIN Technologies, ON, Canada) while single units and LFPs were simultaneously recorded from multi-electrode arrays implanted in the upper limb area of M1 (Supplementary Figure 2a). Monkeys Mk and Ls made movements to one direction (270° and 45°, respectively), and Bx was trained to make movements in two opposing directions (45° and 225°). LFP signals were band-pass filtered centered at the frequency of peak power in the beta frequency range (15-35 Hz: 31±3 Hz for monkeys Mk and Bx, and 21±3 Hz for monkey Ls; see Methods) from which beta amplitude profiles were computed using the Hilbert transform. Traditionally, beta amplitude dynamics are determined by trial-averaging beta profiles in order to reduce noise^25–27^. However, trial-averaging conceals possible interesting beta dynamics that vary from trial to trial^28^. To extract single-trial beta profiles, we used an auto-encoder that was trained to generate de-noised versions of the input beta signals (Supplementary Figure 2b). These single-trial, de-noised profiles exhibited characteristic attenuation (i.e. desynchronization) prior to movement initiation, a mesoscopic signature of cortical excitability^21^. More importantly, these beta dynamics revealed spatial gradients that evolved in time indicative of spatio-temporal patterns that propagated in one of two oppositely oriented directions along a rostro-caudal axis in MI (Figure 1c).

For each trial, we quantified the propagation direction of beta attenuation time prior to movement initiation by first determining when each electrode’s normalized beta profile dropped below a threshold. We then used linear regression to predict beta attenuation times (BATs) from the spatial locations of each electrode (see Equation 1), where the electrode location on the array (*x* and *y* coordinates) was the independent variable and the corresponding BAT was the dependent variable. The regression coefficients could then be used to compute the orientation from early to late BATs (see Equation 2) which we refer to as the beta attenuation orientation (BAO) for that trial. Trials that satisfied the criterion of R^2^ ≥ 0.2 (p ≤ 0.05, F-Statistic; number of trials = 278/366 (76%) for Mk; 1626/1723 (94%) for Ls; 1369/1868 (73%) for Bx making 45 degree arm movements; 1417/1909 (74%) for Bx making 225 degree arm movements where 0, 90, 180 and 270 degree movements correspond to directions to the right, away from the body, left, and into the body, respectively) for the regression fit were retained for further analyses. As suggested by Figure 1c, single trial BAOs were directed in one of two oppositely oriented directions (Figure 1d). For all three monkeys, the distributions of single-trial BAOs (pooled over 4 sessions for monkey Mk, 9 sessions for Ls, 11 sessions for Bx making 45 degree movement and 11 sessions for Bx making 225 degree movement) were bimodally distributed along a roughly rostro-caudal axis (rostromedial-caudolateral axis for Mk and Bx and rostrolateral-caudomedial axis for Ls) albeit the two modes were not always equal in size (Figure 1e). The BAO distributions were well fit by a mixture of von Mises functions model with means of 137° and −20° for Mk; 33° and −126° for Ls; 143° and −33° for Bx for 45 degree arm movements, and 137° and −32° for 225 degree arm movements. The BAO distributions for the two movement directions in Bx were not statistically different (p=0.1, Kuiper’s two sample test). Multi-unit activity was not statistically different for trials in either of the modes and the few trials that fell outside of the two modes (one-way ANOVA for 3 groups, p-values ranging from 0.21 to 0.99 for different datasets; Supplementary Figure 3). However, trials in which the BAO propagated in the rostro-to-caudal direction had significantly shorter reaction times as compared to trials with caudal-to-rostral propagation in two of the monkeys (mean reaction time of 387 and 405 milliseconds, respectively, for Monkey Bx making 45 degree movements, p = 0.0049, two-sample t-test (not significant for 225 degree movements, p=0.4); 119 and 143 milliseconds for Monkey Ls, p = 0.0002, two-sample t-test).

### Propagating sequences and somatotopy

We examined the spatial relationship between the propagating sequences and the somatotopic organization of M1 using suprathreshold electrical stimulation. We documented muscle twitches through visual observation and muscle palpation in our three animals by stimulating each electrode (Supplementary Figure 4). Given the nature of the behavioral task (i.e. a reaching task involving primarily proximal muscles), we confined our analysis of propagating sequences on arrays placed primarily in proximal zone even though we had an additional array (in Ls and Bx) placed in the distal zone. Despite the crude somatotopy as others have documented^29–32^, we found that the axis of propagation was nearly orthogonal to the medio-lateral gradient from proximal to distal representations.

### Causal evidence linking propagating patterns to movement initiation

To provide more direct causal evidence that propagating patterns facilitate movement initiation, we conducted experiments designed to perturb movement initiation in which subthreshold, patterned electrical stimulation across a set of electrodes was delivered during the reaction time period (Figure 2a). We hypothesized that patterned electrical stimulation that propagated against (INCONGRUENT) the natural propagating pattern of beta attenuation would delay movement initiation as compared to patterned stimulation that propagated with (CONGRUENT) the natural propagating pattern (see Methods). Given that the BAO varies from trial to trial (i.e. along the rostro-caudal or caudo-rostral directions), we could not assume a particular BAO on any given stimulation trial. Moreover, stimulation artifacts made it difficult to measure the BAO on stimulation trials. This was not an issue for monkey Ls as the bimodal distribution of BAOs was highly asymmetric on non-stimulation trials (i.e. nearly unimodal). Therefore, we could assume a single BAO direction across stimulation trials (2086 trials recorded over 13 sessions). For monkey Bx where the BAO distribution was nearly symmetric, we devised a method for selecting a subset of stimulation trials that we could assume had a highly unimodal BAO distribution based on no stimulation (NOSTIM) trials (see Methods). Only this subset of trials was analyzed for Bx (657/1772 or 37% of stimulation trials for 45 degree movement and 724/1686 or 43% of stimulation trials for 225 degree movement). Likewise, for monkey Mk, a similar procedure was applied to select a subset of trials (84/187 or 45% of stimulation trials) from one of the datasets that showed a bimodal distribution. The second data set from Mk exhibited a highly unimodal distribution on NOSTIM trials. It should be emphasized that the assumption of unimodality is conservative and likely leads to weaker differential effects on reaction time given that some stimulation trials may have had BAOs directed in the opposite direction.

**Figure 2.**
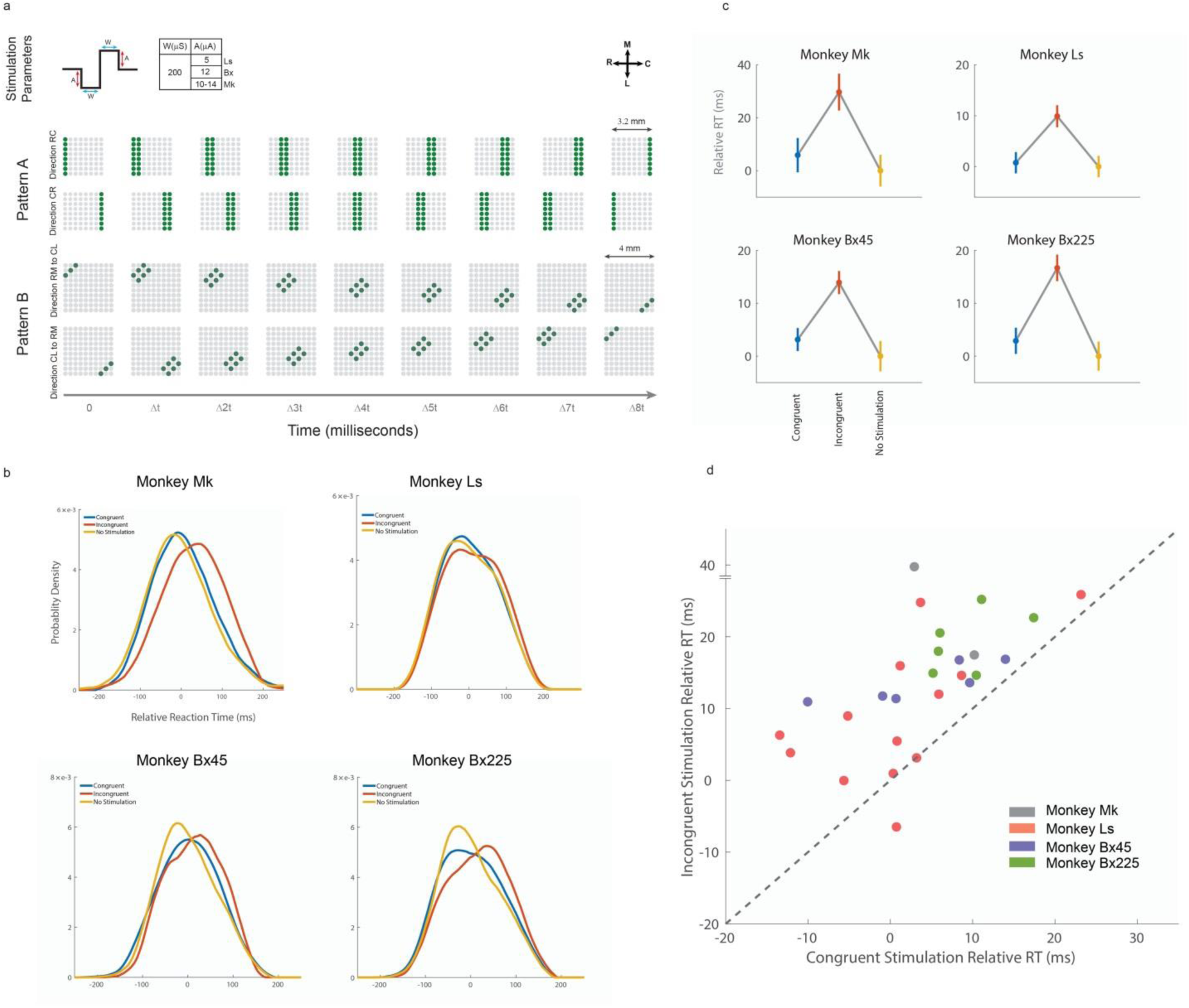
Effect of spatio-temporal stimulation on reaction time. **a.** The stimulus was a biphasic pulse with sub-threshold currents (top), delivered concurrently over sets of electrodes with propagation latency (Δt) of 10 milliseconds (4 milliseconds for Monkey Mk). Stimulation patterns were delivered to a group of electrodes propagating from rostral to causal sites or vice versa (Pattern A, middle) or diagonally (Pattern B, bottom). The stimulus patterns were delivered either along the BAO direction (CONGRUENT) or against the BAO direction (INCONGRUENT). **b.** The distributions of reaction times for the two stimulation conditions along with those trials that received no stimulation (NO STIMULATION) are shown for all three monkeys. **c.** Mean (standard error) relative reaction time in the three conditions. Relative reaction time was computed by subtracting the mean reaction time in the NO STIMULATION condition for each recording session. **d.** Scatter plot comparing the mean reaction times between the two stimulation conditions from 27 individual experimental sessions over all three monkeys. The diagonal dotted line represents the identity line.

Reaction time distributions for the INCONGRUENT condition were significantly delayed as compared to those in the CONGRUENT condition for all three monkeys (Figure 2b; p=0.03 for Mk; p = 0.019 for Ls; p = 0.009 for Bx for 45 degree reaches and p = 0.001 for Bx for 225 degree reaches, two-sample KS test comparing INCONGRUENT and CONGRUENT conditions). No significant difference in RT distributions was observed between CONGRUENT and NO STIM conditions (p=0.49 for Mk; p = 0.64 for Ls; p = 0.17 for Bx for 45 degree reaches and p = 0.08 for Bx for 225 degree reaches, two-sample KS test). Mean reaction times were significantly longer for INCONGRUENT trials as compared to CONGRUENT and NOSTIM trials for all three monkeys (Figure 2c, p < 0.05, F-statistic ANOVA; post-hoc Bonferroni correction). Mean (standard error) reaction times for Mk were 366 (6.5), 390 (7.2) and 357 (6.0) milliseconds for CONGRUENT, INCONGRUENT, and NOSTIM, respectively. Likewise, for Ls, the corresponding mean reaction times were 160 (2.12), 169 (2.18), and 161 (2.15) milliseconds. Monkey Ls tended to anticipate the presentation of the GO cue leading to very short reaction times. For Bx (45 degree reaches) mean reaction times were 375 (2.2), 386 (2.2), 372 (2.9) milliseconds and for Bx (225 degree reaches) they were 363 (2.5), 377 (2.5), 360 (2.75) milliseconds. By comparing session mean reaction times over all 27 stimulation sessions and over all three monkeys, INCONGRUENT session reaction times were significantly longer than those in CONGRUENT sessions (p = 2.4e-5, paired, two-tailed t-test, Figure 2d).

### Functional connectivity emerges along the rostro-caudal axis at movement initiation

If beta attenuation as measured by the LFP is truly a reflection of local network excitability and ultimately activation, we would expect a similar spatio-temporal pattern to emerge among unit activity at movement initiation. To characterize spatio-temporal patterning in neuronal spiking, we estimated functional connectivity among unit activity at different epochs during the reaching behavior in monkeys Bx and Ls (Figure 3a). Monkey Mk was excluded from this analysis due to an inadequate number of channels with spiking activity. We used a Granger-type statistical model applied to point processes^33,34^ to predict multi-unit activity (MUA) from one electrode based on the past MUAs of other electrodes (i.e. source) to infer the directed network connectivity among all simultaneously recorded electrodes (See Methods). The angle of each significant, directed connection between two electrodes was determined based on the spatial locations of the two electrodes. Each significant connection was then binned to create polar histograms (accounting for both excitatory and inhibitory connections as determined by the sign of the sum of history coefficients associated with the source electrode) representing different directions across the motor cortical sheet. During the movement preparation epoch (an epoch that spanned the GO cue so spiking reflected in part generic preparation without movement direction information), polar distributions of functional connectivity were estimated and were fitted by a mixture of two von Mises functions with means of −92° and 100° in Bx for 45 degree reaches (Figure 3b, top) and - 72° and 95° for 225 degree reaches (Figure 3b, middle), and −67° and 128° in Ls for 45 degree reaches (Figure 3b, bottom). For the movement initiation epoch, the von Mises means shifted orientation to −38° and 119° in Bx for 45 degree reaches (Figure 3c, top) and −50° and 125° for 225 degree reaches (Figure 3c, middle), and −123° and 60^0^ in Ls for 45 degree reaches (Figure 3c, bottom). A bimodal distribution (p = 0.01 for Ls, p = 0.004 for Bx 45 degree reaches and p = 3.9e-5 for Bx 225 degree reaches; Rayleigh test for uniform circular distribution modified to test for bimodality) of connectivity emerged along a rostro-caudal axis during movement initiation that was similar to the BAO distribution (p = 0.1 in Bx for 45 degree reaches and p = 0.05 for 225 degree reaches, and p = 0.055 for Ls; Kuiper’s two sample test). Moreover, the functional connectivity distributions for the two movement directions in Bx were not statistically different (p=0.9, Kuiper’s two sample test). In contrast, functional connectivity during movement preparation showed a distribution that was significantly different from the BAO distribution (p = 0.02 in Bx for 45 degree reaches and p = 0.001 for the 225 degree reaches, and p = 0.0001 in Ls; Kuiper’s two sample test).

**Figure 3.**
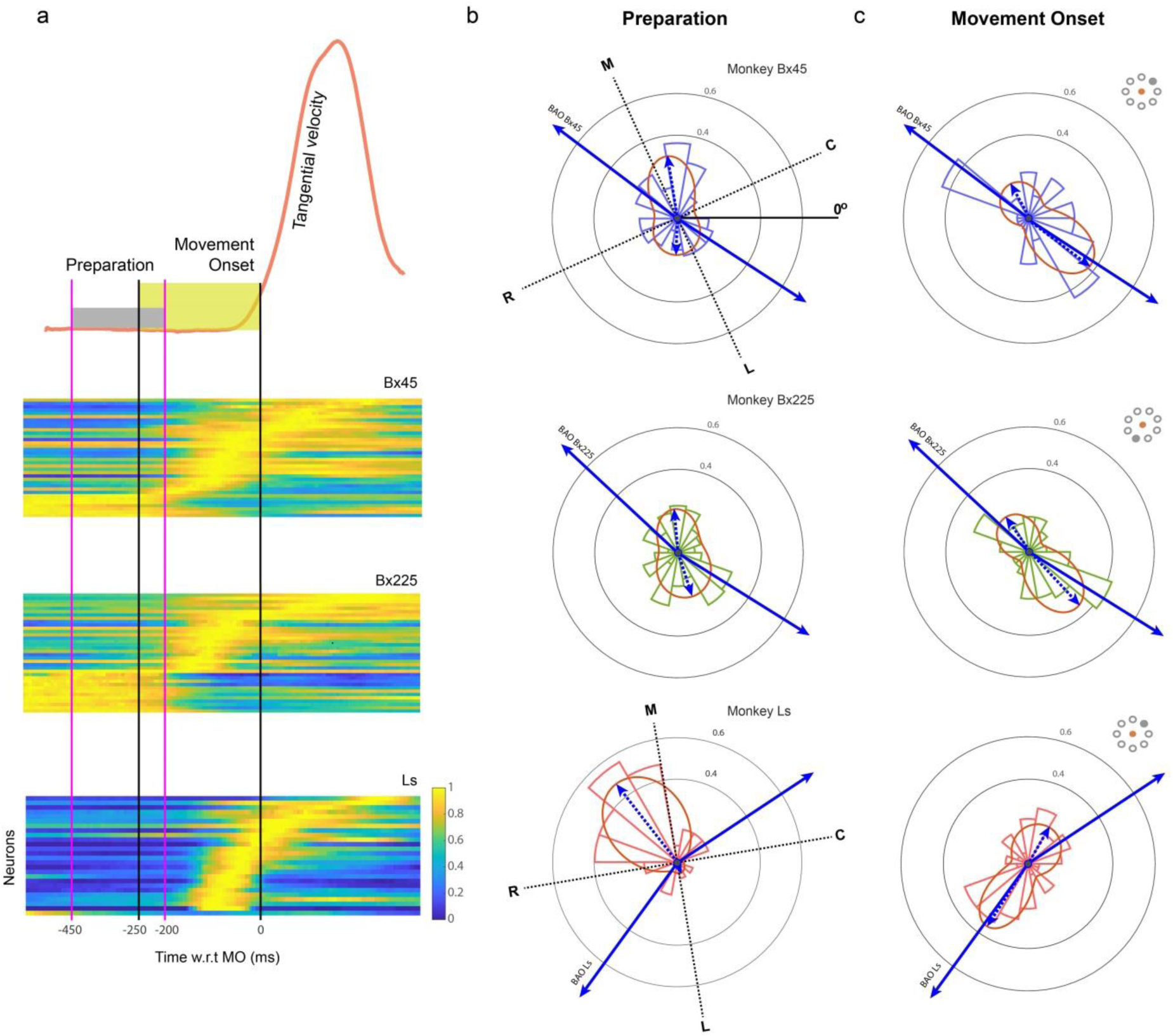
Functional connections among neurons emerge along the BAO axis during movement onset. **a.** Multi-unit activity (MUA) from a population of M1 neurons was used to estimate functional connectivity during movement preparation and movement onset epochs. MUA for each electrode was normalized to its peak activity and sorted by time of peak activity. **b.** Polar histograms of the orientation of functional connections during the movement preparation epoch (top, Bx for 45 degree movements; middle, Bx for 225 degree movements; bottom, Ls for 45 degree movements). The von Mises fits (red curves and dashed blue arrows) show functional connectivity distributions that were different from the BAO distributions. **c.** During the movement onset epoch, the neuronal functional connection distributions were oriented similar to the BAO distributions.

In order to more effectively isolate the neural dynamics of movement planning from movement execution, we also examined functional connectivity during an instructed delay task for monkey Bx where the target stimulus was presented for a random duration ranging from 500 to 1200 milliseconds before a go cue to initiate the movement. Only trials with instructed delay durations longer than 1000 ms were used for the functional connectivity analysis. It is known that single units in primary motor cortex modulate during the instructed delay period where there is no movement^35,36^. We assessed the orientation of directed functional connections during the early (Supplementary Figure 5, left) and late planning (Supplementary Figure 5, right) epochs during the instructed delay period. Mixture models fit to these epochs showed von Mises means that were oriented nearly orthogonal to the BAO axis.

### Propagating patterns along the rostro-caudal axis more reliably predict muscle activity

We hypothesized that downstream areas such as the spinal cord (and motor neuron pools) may be sensitive to the spatio-temporal recruitment order of cortical neurons. Beta oscillation dynamics are likely not involved in the detailed components of movement execution^25^. However, evidence has shown that cortical oscillations and muscle EMGs are related and exhibit significant coherence in the beta frequency range under certain behavioral conditions^37^. More importantly, this coherence can modulate continuously and is related to motor parameters^38^. One way to test the idea that propagating beta attenuation may facilitate muscle recruitment is to decode muscle activity from beta amplitude profiles in cortex and then apply different spatial perturbations to the beta profiles. Using cortical LFP and EMG data from Bx and Ls, we trained a feed-forward neural network (Figure 4a, see Methods) using the Levenberg-Marquardt algorithm^39,40^ to reconstruct a subset of 5 EMG signals (muscles recruited during the task condition; reaching movements to the target at 45°) from a set of cortical beta profiles (Figure 4b and Supplementary Figure 6a). Reconstruction performance on test data not used to train the network was assessed by the fraction of variance accounted for (FVaF in percentage) and ranged from 18% to 55% on all five muscles with an average of 40.5% for Bx and ranged from 15% to 51% on all muscles with an average of 36.5% for Ls. Our goal here was not to develop an accurate decoding algorithm for brain-machine interface applications, but rather to adequately reconstruct muscle activity with a model that could then be used to apply desired spatial perturbations. We applied two kinds of spatial perturbations (random swaps) to the model’s input: a swap 1) parallel to or 2) orthogonal to the BAO axis (See Methods; Figure 4c). A total of 5000 swaps were performed for each perturbation type from which a distribution of FVaFs were generated and compared to each other as well as to the unperturbed FVaF (Figure 4d and Supplementary Figure 6b). Only the parallel perturbation would disrupt the natural sequencing of beta attenuation propagation, and, therefore, we predicted that such perturbations would more effectively disrupt EMG output from the neural network model. Mean FVaFs were, in fact, significantly lower for the parallel perturbations as compared to orthogonal perturbations (p = 6.6e-200 for Bx and p = 3.6e-84 for Ls, two sample t-test).

**Figure 4.**
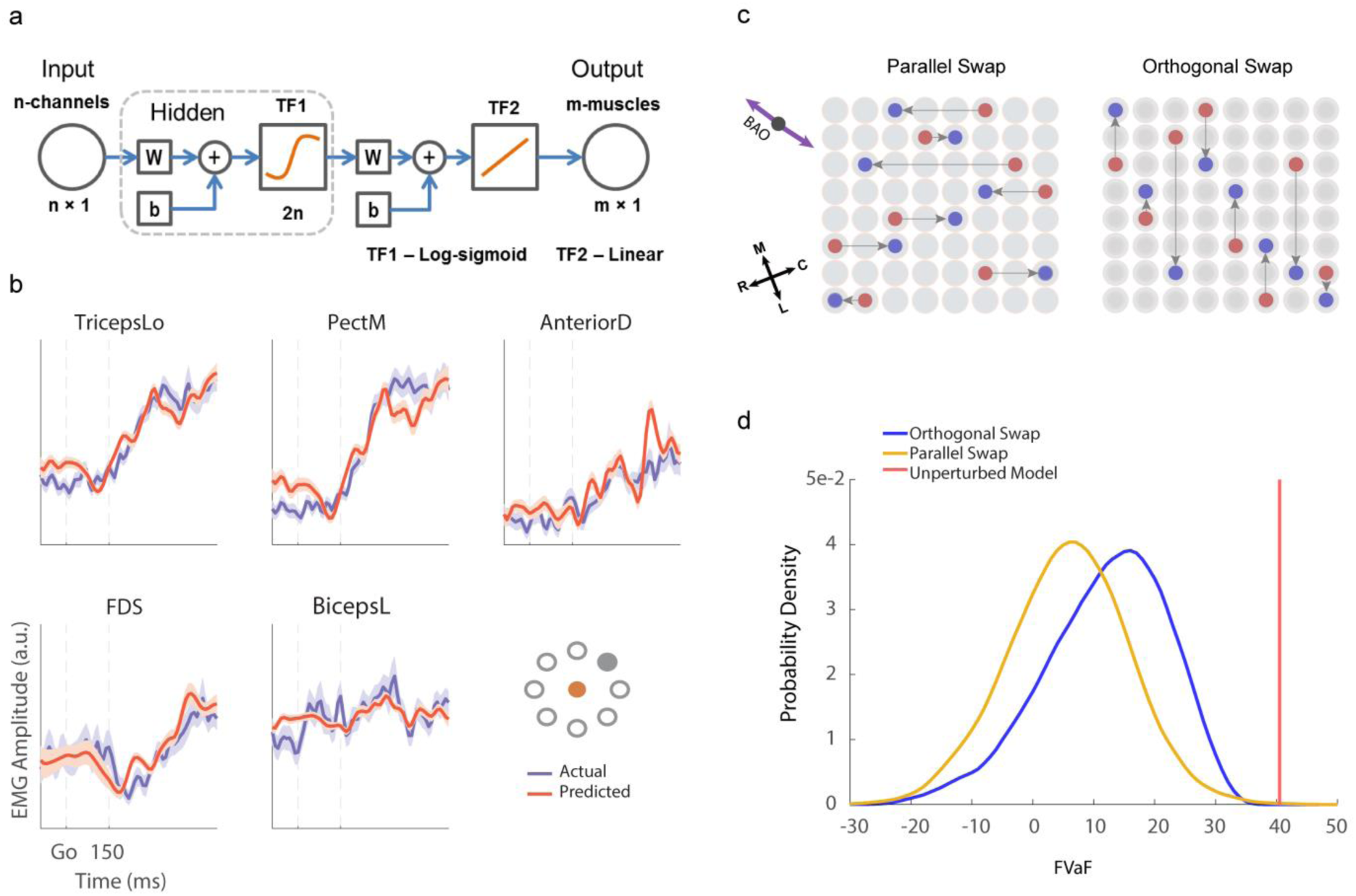
EMG prediction is more sensitive to perturbations parallel to the BAO axis. **a.** A feed-forward back propagation neural network architecture that maps instantaneous beta amplitudes to electromyographic (EMG) from 5 muscles that were active during the particular movement condition. **b.** Trial-averaged predicted EMG values (red) are overlaid on the actual trial-averaged EMG signals (blue) for Bx. **c.** Perturbation of the spatiotemporal pattern was performed along the BAO axis (parallel swap) or orthogonal to it (orthogonal swap), and FVaF of the model’s performance was computed. **d.** Distributions of FVaFs over 5000 swaps are shown for the parallel swap condition (yellow; mean FVaF of 12%) and the orthogonal swap condition (blue; mean FVaF of 6.5%) along with the FVaF of the original unperturbed model (red line; mean FVaF of 40.5%). Inset represent the initial hold (filled red) and target positions (filled gray) of the hand.

## Discussion

This work provides important evidence demonstrating that a specific, propagating rostro-caudal pattern of M1 excitability and unit modulation is required for voluntary movement initiation. First, using subthreshold patterned electrical stimulation, we showed that patterned stimulation that propagates against (INCONGRUENT) the natural propagating direction leads to longer reaction times whereas stimulation propagating with (CONGRUENT) the natural pattern did not affect reaction times. While any form of electrical stimulation will perturb the natural cortical dynamics associated with movement initiation, we suggest that the patterned stimulation propagating against the natural sequence provides a more potent perturbation. Subthreshold ICMS at single sites has been previously shown to lead to increased reaction time in dorsal premotor cortex followed by normal motor execution^41^. Interestingly, similar effects on reaction time were absent for similar single site, subthreshold ICMS in M1. We argue that single site stimulation may not be the ideal paradigm for perturbing the cortical dynamics in M1 associated with movement initiation. Second, functional network connectivity among an ensemble of spiking neurons emerges at movement initiation that is spatially anisotropic and oriented along a similar rostro-caudal axis. This was not observed during movement preparation or planning even though it is known that M1 neurons modulate during these periods (See Figure 3 and Supplementary Figure 5). Spatial structure in functional connectivity was often weak during movement preparation and planning, and, if anything, its spatial anisotropy was oriented orthogonal to the rostro-caudal axis. Anatomical and physiological studies have suggested a spatial anisotropy of horizontal connectivity in M1 that might support these spatially organized propagating sequences. Retrograde axon degeneration following punctate lesions in M1 has indicated longer range horizontal connectivity along the rostro-caudal axis as compared to the medial-lateral axis^42^. Moreover, an ICMS study has provided evidence for dominant functional connectivity along the rostro-caudal axis^43^.

The bimodal distribution of propagation that we observed here is similar in orientation to the bimodal distribution of beta waves (as measured by the phase gradient of the beta oscillations) along the rostro-caudal axis we and others have previously documented to occur during movement preparation^33,44–47^. This may suggest a mechanism by which on-going beta waves propagating along the rostro-caudal axis during movement preparation lead to a sequential pattern of neural activation prior to movement initiation. Beta waves alternately propagate either in the rostral-to-caudal direction or vice versa in a seemingly random fashion during movement preparation. It is also known that excitability is enhanced at particular phases of the beta oscillation^25^ (Murthy and Fetz 1996b). Therefore, as a particular beta wave travels across the cortical sheet in one direction, a wave front of excitability moves across the motor cortex in the same direction. Assuming input originating from outside of motor cortex signaling movement initiation interacts with this beta wave, the combination will more likely lead to firing at that position in motor cortex. The traveling wave front may, therefore, lead to a sequential pattern of activation.

An open question is how these propagation patterns are mechanistically related to appropriate muscle activations for movement initiation. One possibility is that rostro-caudal propagation enhances M1 output through recurrent connectivity due to stronger rostro-caudal horizontal connectivity. However, M1 firing rates for trials when propagating excitability occurred along the rostro-caudal axis were not statistically different from those few trials when propagation occurred outside the rostro-caudal axis. Another possibility is that downstream areas such as the spinal cord are sensitive to the recruitment order of cortical sites. Our muscle decoding results indirectly supports this second alternative. Perturbing the spatial order of beta amplitude inputs along the rostro-caudal axis more effectively disrupts the natural propagating patterns and leads to significantly weaker EMG predictions as compared to orthogonal perturbations which preserve the overall propagation pattern.

The sensitivity of downstream receiving populations to these cortical propagating sequences could be due to the specific characteristics of these patterns. For example, the propagation axis can be thought of as a composition of two phenomena: simultaneous activation of populations orthogonal to the propagating axis (i.e. along the medio-lateral axis) and sequential activation of populations along the propagating axis. Either or both of these phenomena could be responsible for downstream activations that lead to movement initiation. One possibility is that synchronous activation along the medio-lateral axis and rostro-caudal cortical sequencing lead to non-linear amplification of muscle activity which support movement initiation and generation. Further experiments will be needed to shed light on the exact nature of the downstream sensitivity to these natural spatiotemporal patterns.

Our results suggest that these propagating sequences serve a generic role in initiating movements and do not depend on the nature of the movement or the temporal sequencing of limb segments. Although we have not examined this question in great depth, our findings from monkey Bx demonstate that the bimodal distributions of BAOs and functional connectivity are not statistically different between movements toward and away from the animal. Moreover, the spatial structure of these propagating patterns are nearly orthogonal to the medio-lateral gradient from proximal to distal representations. This suggests that these propagating sequences are likely not the result of proximal-to-distal sequencing that is observed in certain motor behaviors^48,49^.

These results have broad significance because they suggest that large-scale, spatially structured propagating patterns of activity may be a common strategy used in various cortical areas to perform behaviorally relevant computations. Spatio-temporal patterns such as traveling planar, radial, circular, and spiral waves have been documented across different cortical areas including somatosensory^50,51^, visual^52–57^, auditory^58,59^ cortices, and hippocampus. Various functional roles for these spatio-temporal patterns have been put forth but it remains a possibility that they are epiphenomena of the recurrently connected cortex^60^. Multi-site, spatio-temporal stimulation using electrical or optogenetic stimulation may be an important approach to provide more direct support for a role of these patterns in cortical processing.

## Acknowledgements

We would like to thank Carrie Anne Balcer for assistance in animal care and training as well as Callum Ross, Juan Alvaro Gallego, and Raeed Chowdhury for assistance in chronic EMG implants. This work was supported by the National Institute of Neurological Disorders and Stroke at the National Institutes of Health (Grant R01 NS045853 to N.G.H.). This work used the Beagle supercomputer resources provided by the Computation Institute and the Biological Sciences Division of the University of Chicago and Argonne National Laboratory, and the Midway cluster at the Research Computing Center of the University of Chicago.

## Author Contributions

K.B., V.P. and W.L. contributed to multiple aspects of the work including surgical procedures, experimental design, data analysis, and manuscript writing/editing as well as data collection for monkeys Ls and Bx. K.B. and N.G.H. were responsible for the single-trial BAO estimation techniques. K.B., K.T. and N.G.H. contributed to the development of the functional connectivity analyses. K.T., M.B. and A.J.S. helped conceive of the patterned stimulation paradigm and contributed to surgical procedures, manuscript editing, and data collection for monkey Mk. N.G.H. supervised the study and contributed to surgical procedures, experimental design, data analysis, and manuscript writing.

## Author information

N.G.H. serves as a consultant for BlackRock Microsystems, Inc., the company that sells the multi-electrode arrays and acquisition system used in this study

Correspondence and requests for materials should be addressed to N.G.H.

## Methods

### Electrophysiology

Three Rhesus (*Macaca Mulatta*) monkeys (Mk, Ls and Bx) were implanted with Utah multi-electrode arrays (1.0 mm electrode length for Mk and Ls; 1.5 mm for Bx and 400 µm pitch from Blackrock Microsystems, Inc. Salt Lake City, UT) in the upper limb region of primary motor cortex (M1), on the contralateral side of the active limb used in the task. Monkeys Ls and Bx received two arrays with 64 recording electrodes per array (8 × 8 grid), positioned on the precentral gyrus along the medio-lateral axis and monkey Mk was implanted with one array of 100 electrodes (10 × 10 grid) (see Supplementary Figure 2a). Surface electrical stimulation during surgical implantation was performed to guide the placement of each electrode array in the arm/hand area of primary motor cortex. Monkeys Ls and Bx were implanted with bi-polar electromyographic (EMG) electrodes in 13 individual muscles (*Anterior Deltoid, Posterior Deltoid, Pectoralis Major, Biceps Lateral, Biceps Medial, Triceps Long head, Triceps Lateral, Brachioradialis, Flexor Carpi Ulnaris, Flexor Digitorum Superficialis, Extensor Carpi Radialis, Extensor Digitorum Communis, Extensor Carpi Ulnaris*). The surgical and behavioral procedures involved in this study were approved by the University of Chicago Institutional Animal Care and Use Committee and conform to the principles outlined in the Guide for the Care and Use of Laboratory Animals.

Neural data recorded as analog signals were amplified with a gain of 5000, bandpass filtered between 0.3Hz and 7.5 kHz, and digitized at 30 kHz. For monkey Mk, the local field potentials were sampled at 2 kHz from the digitized data after low-pass filtering the signals with a 4^th^ order Butterworth filter and a cut-off frequency of 500 Hz. For Ls and Bx, the neural data were zero-phase low-pass filtered with a 10^th^ order Butterworth filter and a cut-off frequency of 500 Hz, and down-sampled to 2 kHz. Multi-unit activity (MUA) was extracted as threshold crossings from the raw data after high-pass filtering with a cut-off frequency of 250 Hz. The threshold limit was set to be −5.5 times of the signal RMS value. For EMG recordings, signal from the thirteen muscles were amplified individually and bandpass filtered between 0.3 and 1 kHz prior to digitization and sampled at 10 kHz.

### Behavioral Task

The animals were operantly trained to perform a center-out planar reaching task using a two-link exoskeletal robot (BKIN Technologies). The tool-tip of the robot determined the position of a cursor on a screen, and the monkeys were able to freely move the robot in the horizontal plane using their upper limb. Monkey Mk used a horizontal screen with his arm and robot underneath the screen, while monkeys Ls and Bx had a vertical screen facing them such that the horizontal movements forward and back corresponded to upward and downward cursor movements. The animals had to hold the cursor on a center target for a fixed duration (600 ms for Mk, 900 ms for Ls, and 1000 ms for Bx) to trigger a peripheral target appearance (GO cue). The animal had to move the cursor to reach the peripheral target to get a juice reward. Monkey Mk was presented with targets at 270° on a circle of radius 6 cm (where 0° corresponded to rightward movements and 180° corresponded to leftward movements), and for monkey Ls the peripheral target was at 45^0^. Monkey Bx was trained with two targets, one at 45° and another at 225°.

### Beta Band and Single-trial Beta Attenuation Orientation

The frequency associated with the peak power in the beta frequency range (15-35 Hz) was estimated from the channel-averaged power spectral density of the neural signals from each array over a large number of trials. For monkey Ls, the beta peak occurred at 21 Hz (approximated to the nearest integer value), and for monkeys Bx and Mk the peak occurred at 31 Hz. For the latter two monkeys, a second smaller peak also occurred at 21 Hz but was not used for further analysis. The signal was then band-pass filtered with a bandwidth of 6 Hz centered on the peak beta, i.e., 21±3 Hz for monkey Ls and 31±3 Hz for monkeys Bx and Mk. The beta envelopes of the signal were computed using the magnitude of the Hilbert transformation applied to the band-pass filtered signal (see Figure 1a).

Single-trial beta envelopes were extracted using an auto-encoder neural network where the inputs are the same as the targets. Segments of the signal (−500 ms to 600 ms with respect to the GO cue) from all the recorded channels were extracted for each trial and decimated to 100 Hz. A vector of length nChannels × mSamples was created for each trial and fed through the auto-encoder (see Supplementary Figure 2b). The hidden layer was set to be half the size of the input layer so as to form a bottleneck architecture, and the data were projected to the output layer. With training, the auto-encoder was able to reconstruct the signal that preserved both spatial patterns and significant temporal variance of the signal. The output beta envelopes from the auto-encoder were then low-pass filtered with a 10^th^ order filter and cut-off frequency of 5 Hz. The filtered signals were normalized to have an amplitude range of [0, 1].

The normalized single-trial beta envelope attenuates and crosses an amplitude threshold value at a particular time (referred to as beta attenuation time or BAT) which varies depending on the channel (see Figure 1d). These BATs, when arranged spatially on their corresponding electrode locations on the array, exhibit a propagating pattern with a particular orientation. The orientation of the beta attenuation (BAO) was estimated using a linear regression fit to the BATs as follows,

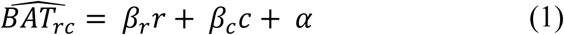

where, *β*_r_ and *β*_c_ are the coefficients of the rows and columns of the array, and *α* denotes the constant offset time. The direction of BAO was then estimated as,

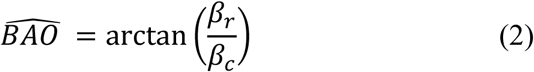

The F-statistic of the model was used to ascertain statistical significance of the fit. For each trial, a BAO and its associated coefficient of determination (R^2^) were calculated. On any given trial, the amplitude threshold for determining the BATs was determined by varying the amplitude from 0.5 to 0.15 in steps of 0.05 and finding the threshold with the largest associated R^2^.

### Patterned Electrical Stimulation

A spatio-temporal pattern of intracortical microstimulation (ICMS) was delivered to a fixed set of electrodes on the single array in Mk and the medial arrays for Ls and Bx with a fixed onset latency after the GO cue (150 ms for Mk, 50 ms for Ls and 200 ms for Bx). The patterned ICMS propagated either rostro-caudally or caudo-rostrally which was either congruent or incongruent to the BAO for a given trial (see Figure 2a). As a control, there were also trials without stimulation. These three conditions (congruent, incongruent, and no stimulation) were randomly interleaved across trials for monkeys Ls and Bx. For monkey Mk, each of the three conditions was presented in multiple 25-trial blocks where each block was randomly given. The patterned ICMS consisted of trains of biphasic (200 µs negative and positive pulses with an interphase latency of 55 µs) pulses with constant subthreshold amplitude (5 µA for Ls and 12 µA for Bx). The subthreshold current for Mk during the experiments was set to be 2 µA lesser than the current levels that resulted in evoked responses during the two stimulation conditions, tested immediately prior to the experimental session. For one data set, the current per electrode was set as 13 µA for the congruent stimulation and 14 µA for the incongruent condition while in the other stimulation data set, the currents were 12 µA and 14 µA for the congruent and incongruent conditions, respectively. For monkeys Ls and Bx, a set of 8 or 16 electrodes were stimulated simultaneously at any given time (see Pattern A in Figure 2a). Pulse trains on a successive column were initiated at a latency of 10 milliseconds relative to the previous column. For Mk, 3 or 6 electrodes were stimulated at any instance, with inter-stimulation latency of 4 milliseconds (see Pattern B in Figure 2a).

### Determining BAO for Trials with Stimulation

For trials that received patterned stimulation, recovering the complete beta envelopes was not possible due to large saturating electrical artifacts during stimulation. For monkey Ls the bimodal BAO distribution was highly asymmetric on non-stimulation trials (i.e. nearly unimodal). Also, for one data session examined in Mk, the BAO distribution was highly unimodal. Therefore, we could assume a single BAO direction across stimulation trials. To determine the BAOs for the stimulation trials in monkey Bx and for the second session in Mk, an alternative approach was taken by examining the dynamics of the beta envelopes in trials when no stimulation occurred. Typically, the channel-averaged beta envelope exhibits a local peak in amplitude at different times prior to attenuation depending on the trial. The approach was to find a time window around the GO cue where the local beta peaks occurred on a subset of no stimulation trials such that their BAO distribution was highly unimodal. This time window (set to 300 ms in duration; 150 ms for monkey Mk) was found by sliding the window in 100 ms steps starting at −300 ms with respect to the GO cue, selecting no stimulation trials whose local beta peak occurred in that window, and computing the BAO distribution (Supplementary Figure 7). The window whose selected trials were highly unimodal (where at least two thirds of the trials in the window were oriented along one BAO mode) in their BAO was chosen and then used to select stimulation trials whose local beta peak occurred in the same window. The assumption was that these stimulation trials exhibited a similar unimodal BAO distribution and, therefore, all these stimulation trials had BAOs pointing roughly in the same direction. To detect local beta peaks in stimulation trials, partial beta envelopes (−500 ms to 200 ms with respect to the GO cue; −500 to 25 ms for monkey Mk) that were not corrupted by electrical artifacts were extracted using the auto-encoder. Whenever multiple time windows exhibited unimodal BAOs in the no stimulation trials, the window that was closer to or overlapping the GO cue was chosen. To compute reaction time means and distributions for all three conditions, we excluded trials whose movement onset times occurred before the onset of the stimulation as well as trials with very long reaction times (>500 ms for Mk and Bx and >300 ms for Ls).

### Computing Reaction Times of Single Trials

Tangential velocity trajectories estimated from the Cartesian position of the reaching kinematics data sampled at 2 kHz were used for reaction time computation. From the normalized trajectories of each trial, movement onset times were computed when the velocity crossed 10% of the peak velocity. When there were smaller velocity peaks that exceeded the 10% threshold prior to final velocity peak associated with the actual movement to the target, movement onset was computed using the final threshold crossing.

### Firing rates

Multi-unit firing rates were calculated for each trial in 150 ms temporal windows, shifted over time with steps of 10 ms. Rates were then averaged over trials to obtain the mean firing rate of each channel’s multi-unit. For the heatmaps of Figure 3, mean firing rates were normalized by dividing by the multi unit’s maximum rate. A different normalization procedure was used to compare the firing rates of each mode. First the average baseline activity over all trials (500ms prior to the go cue) was subtracted from each trial’s firing rate. Then, trials were separated into three groups according to BAO mode (rostro-to-caudal BAO, caudo-to-rostral BAO, out of modes) and the average firing rate of each group was calculated. If the multi-unit displayed suppressed modulation, its rates were converted to the additive inverse. Finally, the three average rates were divided by the global maximum, in order to retain possible mode differences. Population response for the three modes was then estimated by averaging over multi-units. To compare the population responses between modes, we used the normalized unit responses at two different epochs: reaction (from go cue to movement onset) and movement (from movement onset to movement end). A one-way AVOVA was done separately at each epoch [factor: BAO mode (three levels), sample size: number of multi units] to determine differences between the population firing rate across BAO modes.

### Directed Functional Connectivity

A granger-type causality estimation technique generalized for point-processes was used to infer directed connectivity among simultaneously recorded neurons. For each candidate multi-unit, the instantaneous spiking was modeled as its conditional intensity function (CIF), *λ(t|H(t))*, where *H(t)*, is the spiking history of all the multi-units in the ensemble up to time *t*. Using a generalized linear model, the log CIF of each multi-unit was modeled in terms of the spiking of its covariate (i.e. source) multi-units, initially with a history of 60 milliseconds binned at 3 millisecond resolution for each source multi-unit. The final model was then found by optimizing the length of the history term using the AIC (Akaike information criterion). To determine the influence of a source multi-unit on a candidate multi-unit, the log-likelihood ratio between models (i) including and (ii) excluding the source multi-unit was computed, and when the model performance decreased significantly (p ≤ 0.05, χ^2^-test, false detection rate corrected for multiple comparison) with the source multi-unit excluded, a directed connectivity from the source multi-unit to the candidate neuron was inferred.

### Decoding Muscle Activity from Beta Envelopes

A feed-forward neural network was developed to map instantaneous beta amplitudes from multiple channels to the EMG activity of a group of muscles. Normalized beta envelopes from 60 channels (100 samples/second) were projected first through the auto-encoder and then fed into the feed-forward neural network to decode 5 muscles based on their relevance to the task and the quality of the signal (Anterior Deltoid, Pectoralis Major, Biceps Lateral, Triceps Long head, Flexor Digitorum Superficialis). The inputs were vectors of 60 × 1 elements, and the outputs were vectors of 5 × 1 elements. The neural network architecture (see Figure 4a) used 120 hidden neurons with logarithmic sigmoid activation function. The output layer that predicts the EMG activities used a linear transfer function. Of the available trials, 80% were used for training and 20 % for testing. The prediction accuracy was computed for each individual muscle as the fraction of variance accounted for (*FVaF*) between the actual EMG and the EMG predicted from the neural network as,

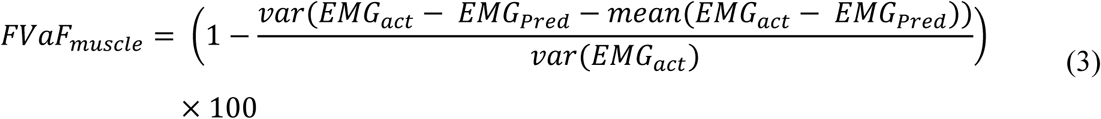

The input to the network was perturbed by swapping the spatial locations of the channels in two different ways. For each swap, beta profiles on eight electrodes were randomly selected on the 64 electrode array and then randomly swapped spatial locations with other beta profiles with the constraint that swapping occurred either parallel to or orthogonal to the BAO axis. A total of 5000 swaps were performed for each of the directions (parallel or orthogonal) and the *FVaF* for each swap was estimated.

### Somatotopic Maps

In order to determine the somatotopy of M1, supra-threshold electrical stimuli (a train of 25 biphasic pulses with 200 µS pulse width per phase and 55 µS interphase interval and 330 Hz; current levels varied from 10 µA to 70 µA per electrode) were delivered to each single electrode one at a time and the corresponding evoked responses documented. Two observers noted the responses while a third individual delivered the stimuli using a Blackrock Microstimulator R96 system (Blackrock Microsystems, Inc. Salt Lake City, UT). Certain muscles were also palpated to identify any evoked responses. We used a 7-point scale (color-coded in Supplementary Figure 4) to label each site from (fingers, red), to wrist (yellow), to elbow (baby blue), to shoulder (blue) joint and/or muscle twitches.

**Supplementary Figure 1.**
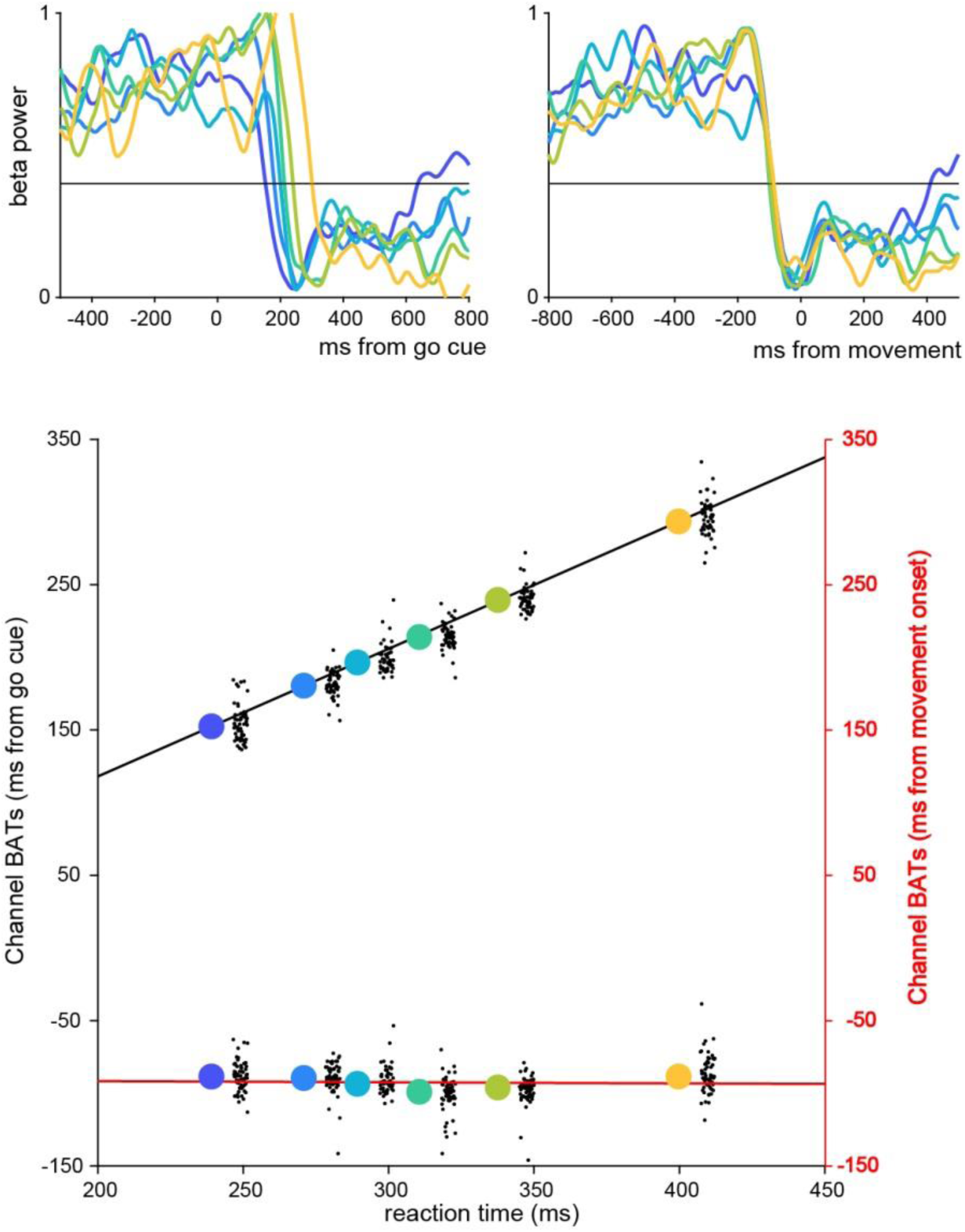
Beta attenuation times are correlated with reaction times and time-locked to movement initiation. Normalized beta envelopes averaged over different trial groups from one session in monkey Bx. Trials were sorted based on reaction time and grouped in bins of 55 trials. Color represents reaction time average of each group from shortest (blue) to longest (yellow) trials. Average beta power for each group is plotted aligned on reaction time (top left) or on movement onset (top right). (bottom) Channel-averaged beta attenuation time regressed on mean reaction time in each trial group with respect to the GO cue (black line, r^2^ = 0.998, p = 8e-9) or movement onset (red line r^2^ = 0.009, p = 0.853). Each big colored dot represents channel-averaged values and each black dot represents single channel values. Color represents reaction time average of each group from fastest (blue) to slowest (yellow) trials.

**Supplementary Figure 2.**
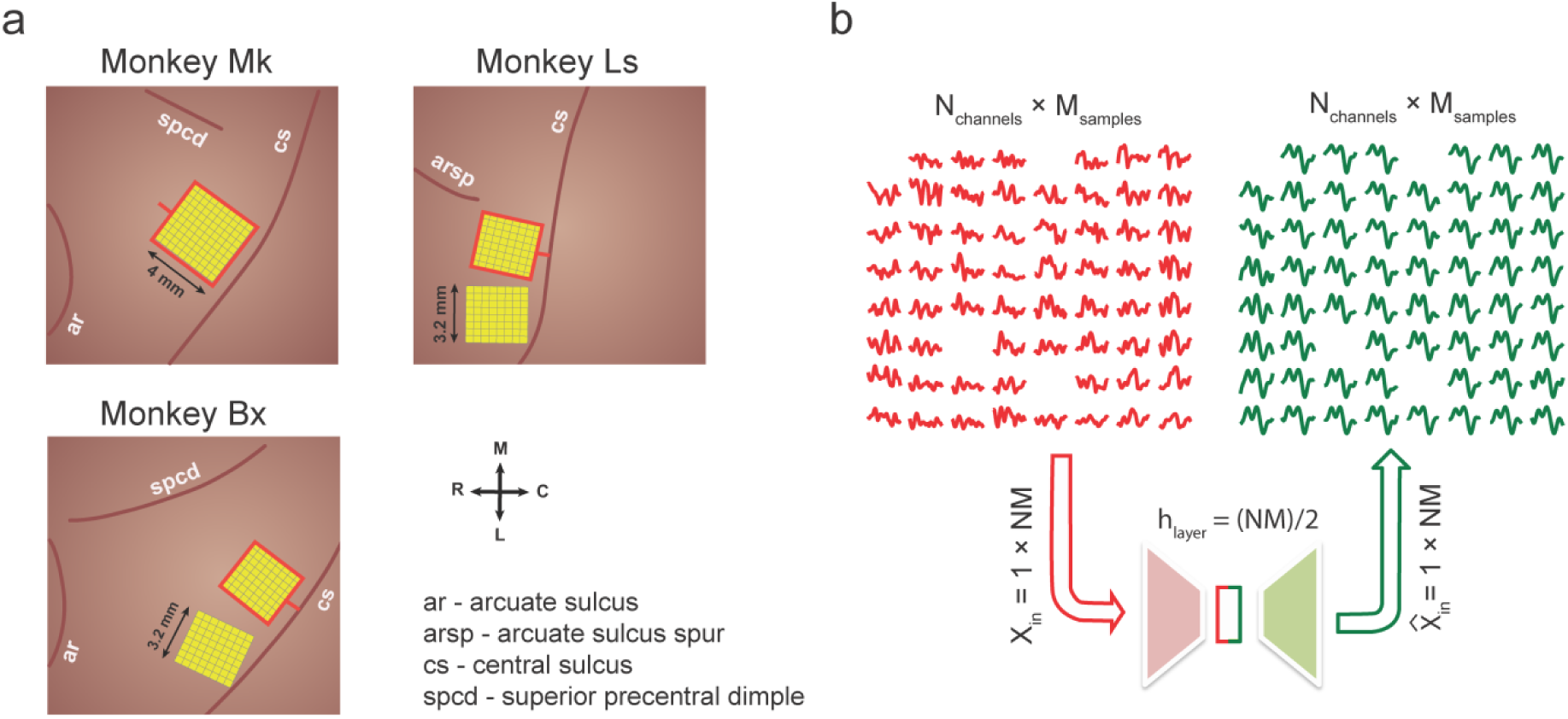
Multi-electrode array implantation and autoencoder architecture. **a.** Multi-electrode arrays were implanted in the primary motor cortex (M1) of three macaques (Mk, Ls and Bx). The arrays marked with red border were used for analyses and patterned stimulation experiments. **b.** Single trial beta envelopes were auto-encoded to de-noise the neural signals. Beta envelopes from N-channels sampled at 100 Hz (M-samples) were projected using the auto-encoder. The hidden layer consisted of (NM)/2 neurons, resembling a bottleneck architecture.

**Supplementary Figure 3.**
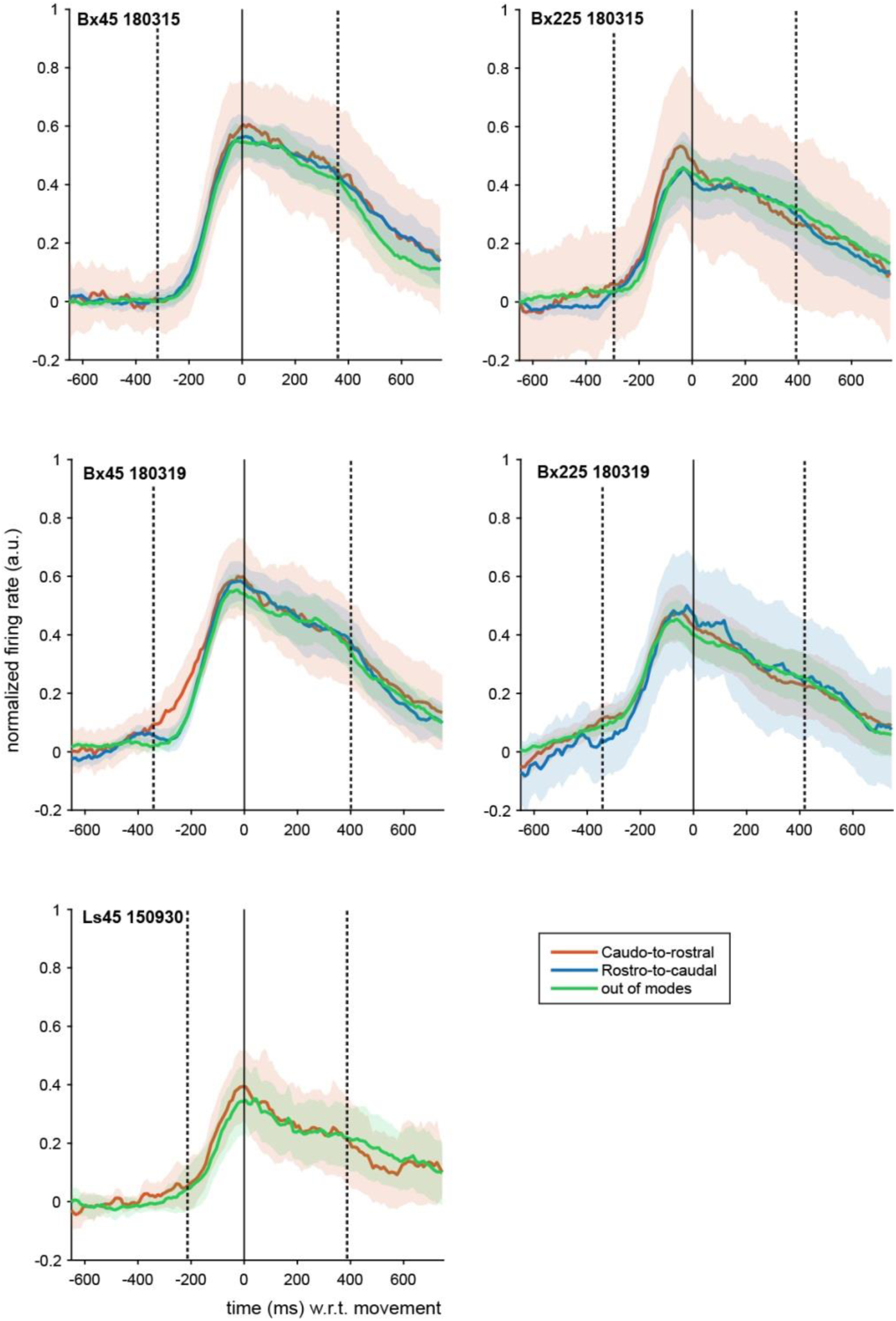
Population firing rate profiles in and outside BAO modes. Population peri-movement time histograms based on 32 to 60 multi-unit sites for trials in the caudal-to-rostral BAO mode (red line), rostral-to-caudal BAO mode (blue line), and trials outside of the two modes (green line) for representative datasets. Trials were aligned on movement onset (black vertical line) and each multi-unit site’s firing rate profile was normalized before averaging (see Methods). Dotted vertical lines represent the median go cue and movement end. Shaded areas represent standard error of the mean.

**Supplementary Figure 4.**
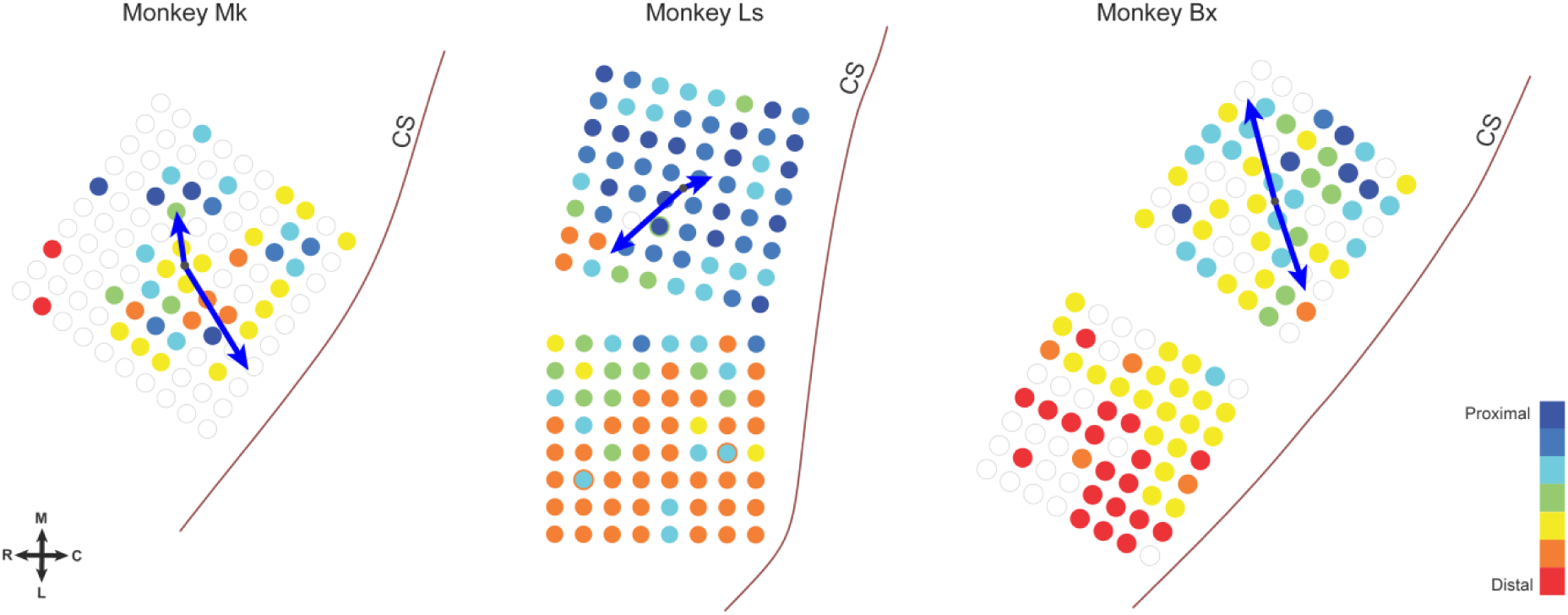
Relationship between propagating sequence directions and somatotopy. The somatotopic maps across the arrays implanted in the three monkeys show the proximal-to-distal somatotopy oriented along the medio-lateral axes. The somatotopic maps for Ls and Bx are based on two 8×8 arrays implanted medio-laterally in M1. The BAO axes (blue arrows), computed for the medial arrays in Ls and Bx, and the one array in Mk, propagate nearly orthogonal to the proximal-to-distal gradient.

**Supplementary Figure 5.**
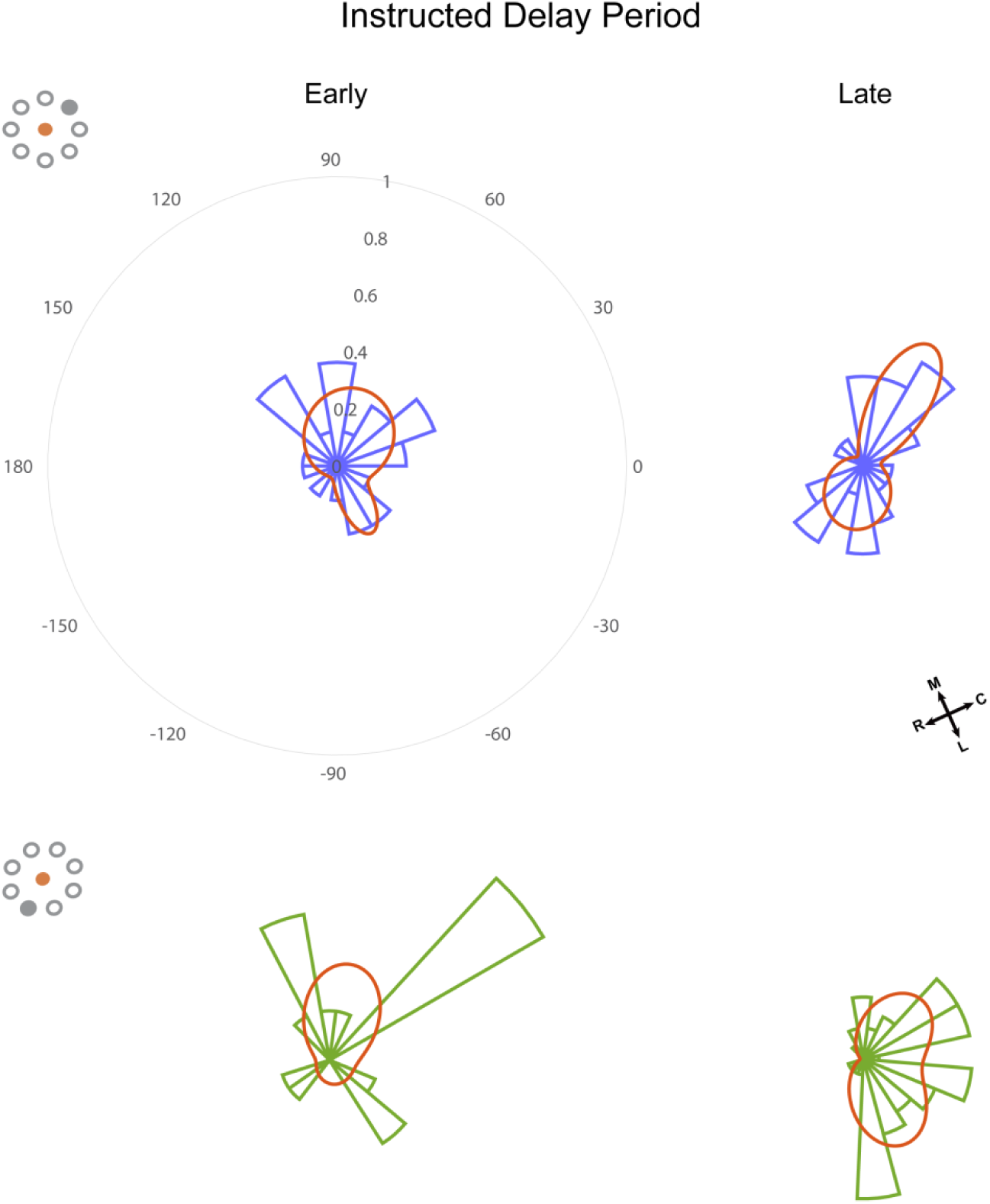
Distribution of Functional Connections during Movement Planning in an Instructed Delay Task. Directional distributions of functional connections estimated using multi-unit activity is shown for early (100 to 350 ms after the instruction cue onset) and late (600 to 850 ms after the instruction cue onset) phases of the instructed delay period. The top panels are for monkey Bx reaching to the target at 45 degrees, and the bottom panels corresponds to reaches at 225 degrees. The distributions were fit with a mixture of two von Mises functions (red curves).

**Supplementary Figure 6.**
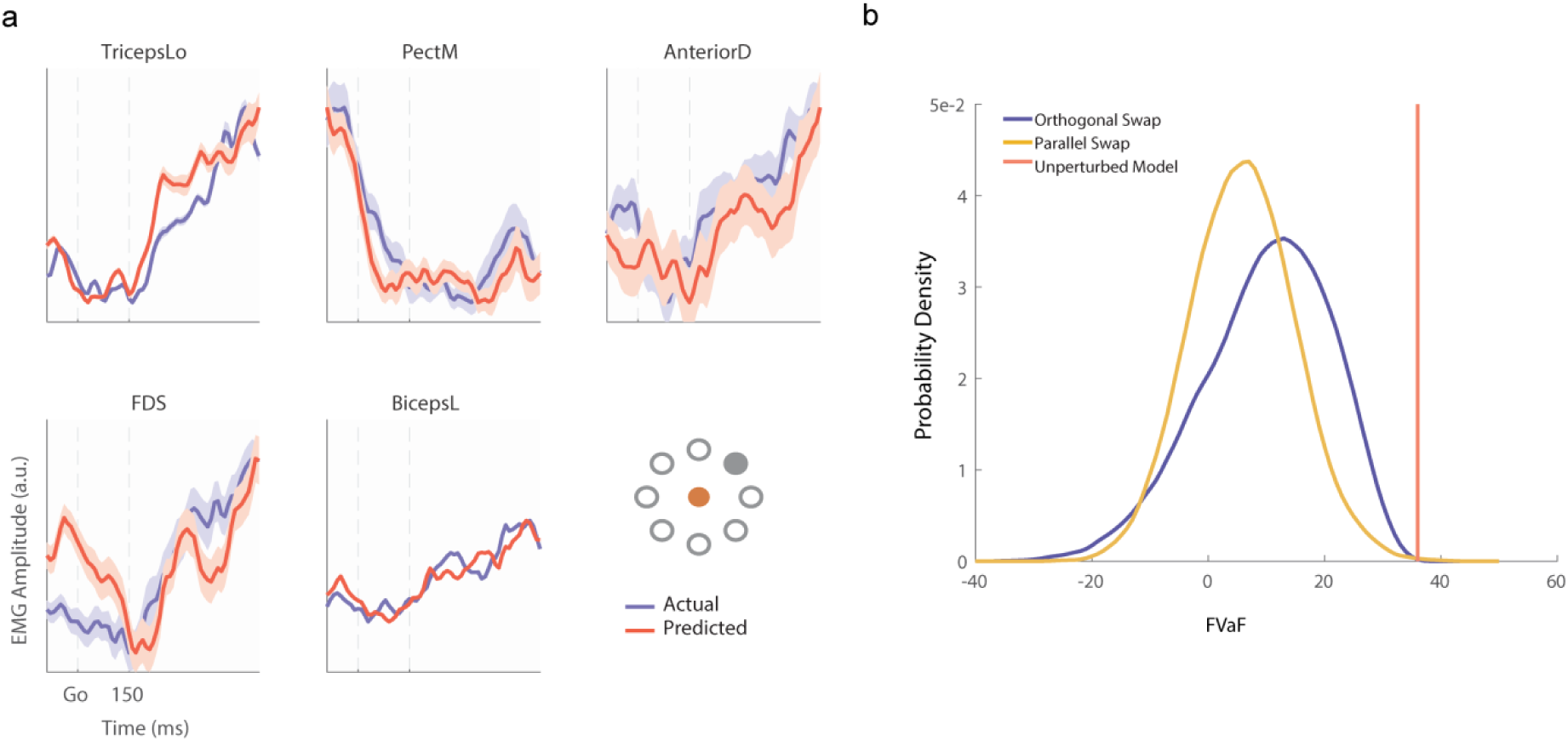
Effects of spatial perturbations on EMG predictions in monkey Ls. **a.** Trial-average predicted EMG (red) and actual EMG (blue) and profiles using a neural network trained with beta envelopes from Monkey Ls. **b.** Distributions of FVaFs over 5000 swaps are shown for the parallel swap condition (yellow; mean FVaF of 5%) and the orthogonal swap condition (blue; mean FVaF of 10%) along with the FVaF of the original unperturbed model (red line; mean FVaF of 36.5% (varying from 15 % to 51% across muscles). Inset represent the initial hold (filled red) and target positions (filled gray) of the hand.

**Supplementary Figure 7.**
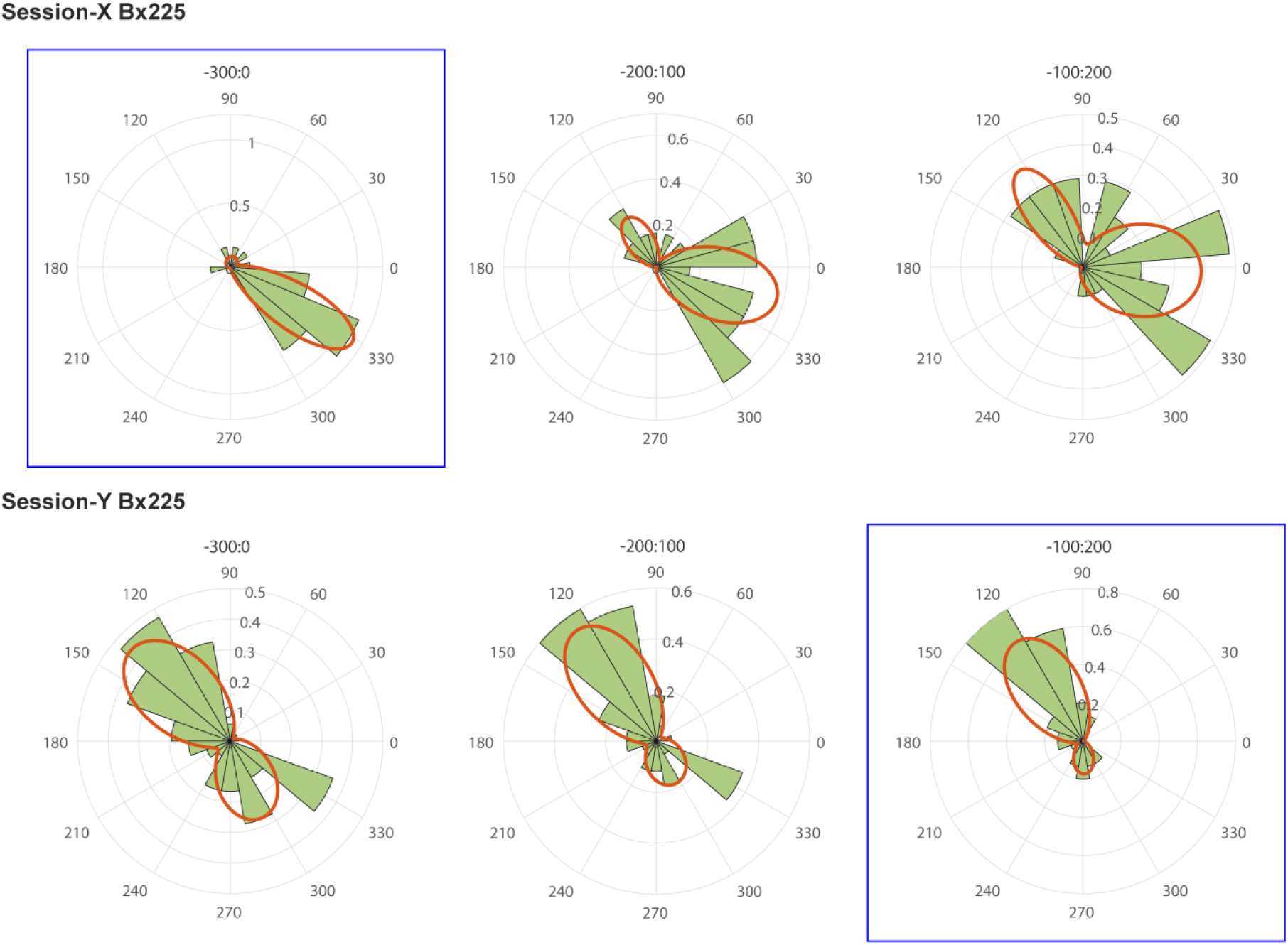
Identification of unimodal BAO window for trial selection. The panels in the two rows (corresponding to two sessions) show the BAO distribution for trials with no stimulation whose channel-averaged beta envelopes had their peaks in the time window [- 300ms, 0ms] (left), [-200ms 100ms] (middle), or [-100ms 200ms] (right) with respect to the GO cue. The window with highly unimodal von Mises fit was selected as the preferred window (top left and bottom right panels highlighted by blue squares), and those trials in the stimulation condition whose peaks fell within this window were assumed to be unimodal in their BAO distribution.

## References

1. Maynard, E. M. et al. Neuronal Interactions Improve Cortical Population Coding of Movement Direction. J. Neurosci. 19, 8083–8093 (1999).

2. Churchland, M. M. et al. Neural population dynamics during reaching. Nature 487, 51–56 (2012).

3. Riehle, A. & Requin, J. The predictive value for performance speed of preparatory changes in neuronal activity of the monkey motor and premotor cortex. Behav. Brain Res. 53, 35–49 (1993).

4. Solodkin, A., Hlustik, P., Chen, E. E. & Small, S. L. Fine Modulation in Network Activation during Motor Execution and Motor Imagery. Cereb. Cortex 14, 1246–1255 (2004).

5. Tkach, D., Reimer, J. & Hatsopoulos, N. G. Congruent Activity during Action and Action Observation in Motor Cortex. J. Neurosci. 27, 13241–13250 (2007).

6. Churchland, M. M., Santhanam, G. & Shenoy, K. V. Preparatory Activity in Premotor and Motor Cortex Reflects the Speed of the Upcoming Reach. J. Neurophysiol. 96, 3130–3146 (2006).

7. Hatsopoulos, N. G. & Suminski, A. J. Sensing with the Motor Cortex. Neuron 72, 477–487 (2011).

8. Hanes, D. P. & Schall, J. D. Neural Control of Voluntary Movement Initiation. Science 274, 427–430 (1996).

9. Vigneswaran, G., Philipp, R., Lemon, R. N. & Kraskov, A. M1 Corticospinal Mirror Neurons and Their Role in Movement Suppression during Action Observation. Curr. Biol. 23, 236–243 (2013).

10. He, S. Q., Dum, R. P. & Strick, P. L. Topographic organization of corticospinal projections from the frontal lobe: motor areas on the lateral surface of the hemisphere. J. Neurosci. 13, 952–980 (1993).

11. Hatsopoulos, N., Joshi, J. & O’Leary, J. G. Decoding Continuous and Discrete Motor Behaviors Using Motor and Premotor Cortical Ensembles. J. Neurophysiol. 92, 1165–1174 (2004).

12. Bullock, D. & Grossberg, S. Neural dynamics of planned arm movements: Emergent invariants and speed-accuracy properties during trajectory formation. Psychol. Rev. 95, 49–90 (1988).

13. Pompeiano, O. Functional Organization of the Cerebellar Projections to the Spinal Cord. in Progress in Brain Research (eds. Fox, C. A. & Snider, R. S.) 25, 282–321 (Elsevier, 1967).

14. Chase, M. H. Synaptic mechanisms and circuitry involved in motoneuron control during sleep. Int Rev Neurobiol 24, 213–258 (1983).

15. Kaufman, M. T., Churchland, M. M., Ryu, S. I. & Shenoy, K. V. Cortical activity in the null space: permitting preparation without movement. Nat. Neurosci. 17, 440–448 (2014).

16. Sanes, J. N., Donoghue, J. P., Thangaraj, V., Edelman, R. R. & Warach, S. Shared neural substrates controlling hand movements in human motor cortex. Science 268, 1775–1777 (1995).

17. Donoghue, J. P., Sanes, J. N., Hatsopoulos, N. G. & Gaál, G. Neural Discharge and Local Field Potential Oscillations in Primate Motor Cortex During Voluntary Movements. J. Neurophysiol. 79, 159–173 (1998).

18. Murthy, V. N. & Fetz, E. E. Coherent 25- to 35-Hz oscillations in the sensorimotor cortex of awake behaving monkeys. Proc. Natl. Acad. Sci. 89, 5670–5674 (1992).

19. Georgopoulos, A. P., Kalaska, J. F., Caminiti, R. & Massey, J. T. On the relations between the direction of two-dimensional arm movements and cell discharge in primate motor cortex. J. Neurosci. 2, 1527–1537 (1982).

20. Chen, R., Yaseen, Z., Cohen, L. G. & Hallett, M. Time course of corticospinal excitability in reaction time and self-paced movements. Ann. Neurol. 44, 317–325 (1998).

21. Schulz, H., Übelacker, T., Keil, J., llerMü, N. & Weisz, N. Now I am Ready—Now I am not: The Influence of Pre-TMS Oscillations and Corticomuscular Coherence on Motor-Evoked Potentials. Cereb. Cortex 24, 1708–1719 (2014).

22. Kühn, A. A. et al. Event-related beta desynchronization in human subthalamic nucleus correlates with motor performance. Brain 127, 735–746 (2004).

23. Williams, D. et al. The relationship between oscillatory activity and motor reaction time in the parkinsonian subthalamic nucleus. Eur. J. Neurosci. 21, 249–258 (2005).

24. Best, M. D., Suminski, A. J., Takahashi, K., Brown, K. A. & Hatsopoulos, N. G. Spatio-Temporal Patterning in Primary Motor Cortex at Movement Onset. Cereb. Cortex 27, (2017).

25. Murthy, V. N. & Fetz, E. E. Oscillatory activity in sensorimotor cortex of awake monkeys: synchronization of local field potentials and relation to behavior. J. Neurophysiol. 76, 3949–3967 (1996).

26. Pfurtscheller, G. & Lopes da Silva, F. H. Event-related EEG/MEG synchronization and desynchronization: basic principles. Clin. Neurophysiol. 110, 1842–1857 (1999).

27. Salmelin, R. & Hari, R. Spatiotemporal characteristics of sensorimotor neuromagnetic rhythms related to thumb movement. Neuroscience 60, 537–550 (1994).

28. Pernet, C. R., Sajda, P. & Rousselet, G. A. Single-Trial Analyses: Why Bother? Front. Psychol. 2, (2011).

29. Kwan, H. C., Mackay, W. A., Murphy, J. T. & Wong, Y. C. An intracortical microstimulation study of output organization in precentral cortex of awake primates. J. Physiol. (Paris) 74, 231–233 (1978).

30. Kwan, H. C., MacKay, W. A., Murphy, J. T. & Wong, Y. C. Spatial organization of precentral cortex in awake primates. II. Motor outputs. J. Neurophysiol. 41, 1120–1131 (1978).

31. Park, M. C., Belhaj-Saïf, A. & Cheney, P. D. Properties of Primary Motor Cortex Output to Forelimb Muscles in Rhesus Macaques. J. Neurophysiol. 92, 2968–2984 (2004).

32. Park, M. C., Belhaj-Saïf, A., Gordon, M. & Cheney, P. D. Consistent Features in the Forelimb Representation of Primary Motor Cortex in Rhesus Macaques. J. Neurosci. 21, 2784–2792 (2001).

33. Takahashi, K. et al. Large-scale spatiotemporal spike patterning consistent with wave propagation in motor cortex. Nat. Commun. 6, 7169 (2015).

34. Kim, S., Putrino, D., Ghosh, S. & Brown, E. N. A Granger Causality Measure for Point Process Models of Ensemble Neural Spiking Activity. PLOS Comput. Biol. 7, e1001110 (2011).

35. Hocherman, S. & Wise, S. P. Effects of hand movement path on motor cortical activity in awake, behaving rhesus monkeys. Exp. Brain Res. 83, 285–302 (1991).

36. Crammond, D. J. & Kalaska, J. F. Prior Information in Motor and Premotor Cortex: Activity During the Delay Period and Effect on Pre-Movement Activity. J. Neurophysiol. 84, 986–1005 (2000).

37. Baker, S. N., Olivier, E. & Lemon, R. N. Coherent oscillations in monkey motor cortex and hand muscle EMG show task-dependent modulation. J. Physiol. 501, 225–241 (1997).

38. Kilner, J. M., Baker, S. N., Salenius, S., Hari, R. & Lemon, R. N. Human Cortical Muscle Coherence Is Directly Related to Specific Motor Parameters. J. Neurosci. 20, 8838–8845 (2000).

39. Hagan, M. T. & Menhaj, M. B. Training feedforward networks with the Marquardt algorithm. IEEE Trans. Neural Netw. 5, 989–993 (1994).

40. Marquardt, D. An Algorithm for Least-Squares Estimation of Nonlinear Parameters. J. Soc. Ind. Appl. Math. 11, 431–441 (1963).

41. Churchland, M. M. & Shenoy, K. V. Delay of Movement Caused by Disruption of Cortical Preparatory Activity. J. Neurophysiol. 97, 348–359 (2007).

42. Gatter, K. C., Sloper, J. J. & Powell, T. P. The intrinsic connections of the cortex of area 4 of the monkey. Brain J. Neurol. 101, 513–541 (1978).

43. Hao, Y., Riehle, A. & Brochier, T. G. Mapping Horizontal Spread of Activity in Monkey Motor Cortex Using Single Pulse Microstimulation. Front. Neural Circuits 10, (2016).

44. Rubino, D., Robbins, K. A. & Hatsopoulos, N. G. Propagating waves mediate information transfer in the motor cortex. Nat. Neurosci. 9, 1549–1557 (2006).

45. Takahashi, K., Saleh, M., Penn, R. D. & Hatsopoulos, N. Propagating Waves in Human Motor Cortex. Front. Hum. Neurosci. 5, (2011).

46. Rule, M. E., Vargas-Irwin, C., Donoghue, J. P. & Truccolo, W. Phase reorganization leads to transient β-LFP spatial wave patterns in motor cortex during steady-state movement preparation. J. Neurophysiol. 119, 2212–2228 (2018).

47. Denker, M. et al. LFP beta amplitude is linked to mesoscopic spatio-temporal phase patterns. Sci. Rep. 8, 5200 (2018).

48. Hirashima, M., Kadota, H., Sakurai, S., Kudo, K. & Ohtsuki, T. Sequential muscle activity and its functional role in the upper extremity and trunk during overarm throwing. J. Sports Sci. 20, 301–310 (2002).

49. Karst, G. M. & Hasan, Z. Timing and magnitude of electromyographic activity for two-joint arm movements in different directions. J. Neurophysiol. 66, 1594–1604 (1991).

50. Petersen, C. C. H., Grinvald, A. & Sakmann, B. Spatiotemporal Dynamics of Sensory Responses in Layer 2/3 of Rat Barrel Cortex Measured In Vivo by Voltage-Sensitive Dye Imaging Combined with Whole-Cell Voltage Recordings and Neuron Reconstructions. J. Neurosci. 23, 1298–1309 (2003).

51. Ferezou, I. et al. Spatiotemporal Dynamics of Cortical Sensorimotor Integration in Behaving Mice. Neuron 56, 907–923 (2007).

52. Sato, T. K., Nauhaus, I. & Carandini, M. Traveling Waves in Visual Cortex. Neuron 75, 218–229 (2012).

53. Zanos, T. P., Mineault, P. J., Nasiotis, K. T., Guitton, D. & Pack, C. C. A Sensorimotor Role for Traveling Waves in Primate Visual Cortex. Neuron 85, 615–627 (2015).

54. Han, F., Caporale, N. & Dan, Y. Reverberation of Recent Visual Experience in Spontaneous Cortical Waves. Neuron 60, 321–327 (2008).

55. Xu, W., Huang, X., Takagaki, K. & Wu, J. Compression and Reflection of Visually Evoked Cortical Waves. Neuron 55, 119–129 (2007).

56. Muller, L., Reynaud, A., Chavane, F. & Destexhe, A. The stimulus-evoked population response in visual cortex of awake monkey is a propagating wave. Nat. Commun. 5, 3675 (2014).

57. Prechtl, J. C., Cohen, L. B., Pesaran, B., Mitra, P. P. & Kleinfeld, D. Visual stimuli induce waves of electrical activity in turtle cortex. Proc. Natl. Acad. Sci. 94, 7621–7626 (1997).

58. Song, W.-J. et al. Cortical Intrinsic Circuits Can Support Activity Propagation through an Isofrequency Strip of the Guinea Pig Primary Auditory Cortex. Cereb. Cortex 16, 718–729 (2006).

59. Witte, R. S., Rousche, P. J. & Kipke, D. R. Fast wave propagation in auditory cortex of an awake cat using a chronic microelectrode array. J. Neural Eng. 4, 68 (2007).

60. Muller, L., Chavane, F., Reynolds, J. & Sejnowski, T. J. Cortical travelling waves: mechanisms and computational principles. Nat. Rev. Neurosci. 19, 255–268 (2018).

